# An Integrative Network-based Analysis Reveals Gene Networks, Biological Mechanisms, and Novel Drug Targets in Alzheimer’s Disease

**DOI:** 10.1101/853580

**Authors:** Zachary F Gerring, Eric R Gamazon, Anthony White, Eske M Derks

## Abstract

Alzheimer’s disease is a highly heritable and severe neuropsychiatric condition. Genome-wide association studies have identified multiple genetic risk factors underlying susceptibility to Alzheimer’s disease, however their functional impact remains poorly understood. To overcome this shortcoming, we integrated genome-wide association summary statistics (71,880 cases, 338,378 controls) with tissue-specific gene co-expression networks derived from GTEx to identify functional gene co-expression networks underlying the disease. We found genetic variants associated with Alzheimer’s disease are enriched in gene co-expression networks involved in immune response-related biological processes. The implicated gene co-expression networks are preserved across multiple brain and peripheral tissues, highlighting the potential utility of peripheral tissues in genetic studies of Alzheimer’s disease. We also performed a computational drug repositioning analysis by integrating gene expression changes within Alzheimer’s disease modules with drug-gene signatures from the Connectivity Map, and show disease implicated networks retrieve known Alzheimer’s disease drugs and novel drug repurposing candidates for follow-up functional studies. Our results improve the biological interpretation of recent genetic data for Alzheimer’s disease and provide a list of potential anti-dementia drug repositioning candidates of which the efficacy should be investigated in functional validation studies.

## Introduction

Alzheimer’s disease (AD) is a common neurodegenerative disorder, characterised in its early stages by mild memory loss and progressing to severe impairment of broad executive and cognitive functions. The most common form of Alzheimer’s disease (late onset Alzheimer’s disease) typically affects those age over 65 years of age and has a complex molecular background, driven in part by a polygenic mode of inheritance. A recent genome-wide association study (GWAS) meta-analysis of 71,880 AD cases and proxy cases and 383,378 controls identified 20 disease-associated loci [1]. Detailed functional studies showed these loci harbour common (minor allele frequency, MAF > 0.01) single nucleotide polymorphisms (SNPs) that regulate the activity of genes in immune-related peripheral tissues (whole blood, liver, and spleen), as well as microglial cells—the chief immune cells of the brain. Furthermore, biological pathway analysis of the implicated genes showed enrichment of previously associated lipid system pathways, highlighting a potential integrated mechanism between dysfunctional lipid metabolism and immune responses in the brain [2].

Genetic risk factors for disease may converge on highly correlated groups of genes that interact with one-another to alter the activity of multiple biological pathways and cellular processes in a disease relevant tissue [3]. Gene expression is an intermediate molecular phenotype that is directly modified by DNA sequence variation (expression quantitative trait loci; eQTLs), epigenetic marks such as DNA methylation, and the environment, as well as the expression of other genes [4]. Gene expression analyses of post-mortem brain tissue have identified distinct cell types and biological pathways underlying AD pathogenesis [5,6]. These studies are largely based on tests of association with individual genes or groups of curated genes with a common biological function. An alternative approach is to study how genes interact with one-another using gene co-expression analysis, which take the correlation between every gene pair expressed in a particular (tissue) sample to generate a molecular substrate for association testing with a disease state [7]. We recently built gene co-expression networks using 48 human tissues and cell types collected for the Genotype-Tissue Expression (GTEx) study [8]. We used these data to test for the enrichment of depression GWAS signals within gene co-expression modules (or groups of highly correlated genes), under the biologically valid assumption that connectivity among genes may be leveraged to identify genes not directly implicated in disease.

Co-expression networks can also be used as a functional substrate for the integration of other types of molecular data for the identification and function of disease-associated genes. This includes new types of chemical libraries that describe the effects of a given drug compound on gene expression, known as a drug-gene database. The Connectivity Map, known as CMap [9], contains gene expression signatures resulting from genetic and pharmacologic perturbagens measured across multiple cell types. Drug-gene signatures—that is, gene expression changes following a genetic or pharmacologic perturbagen—can be integrated with disease-associated gene expression changes to identify compounds that might “normalise” gene expression. Characterising the complex interactions between genes in a network-based framework may identify targets for potential treatments through computational drug repositioning. Therefore, we aim to integrate tissue-specific gene co-expression networks with AD association signals and drug-gene signature data to identify and prioritise drug compounds that target disease processes.

## Methods

### Alzheimer’s disease GWAS summary statistics

Detailed methods, including a description of population cohorts, quality control of raw SNP genotype data, and association analyses for the Alzheimer’s disease GWAS is described in detail elsewhere [1]. The Alzheimer’s disease GWAS was performed in a three-stage meta-analysis. The first phase consisted of 24,087 AD cases and 55,058 controls collected by the Alzheimer’s disease working group of the Psychiatric Genomics Consortium (PGC-ALZ), the International Genomics of Alzheimer’s Project (IGAP), and the Alzheimer’s Disease Sequencing Project (ADSP). All cases in phase 1 received clinical confirmation of late-onset Alzheimer’s disease. The second phase included 47,793 proxy cases and 328,320 proxy controls from the UK Biobank (UKBB); proxy cases where defined as individuals with one or both parents diagnosed with AD, while proxy controls were defined as individuals with parents who do not have AD. Phase 3 involved the meta-analysis of phase 1 and phase 2 cohorts, the results of which were tested for replication in an additional independent case-control sample from deCODE (6,593 AD cases and 174,289 controls). Raw genotype data for each cohort were processed according to a standardised quality control pipeline [1,10]. Logistic regression association tests were performed on imputed marker dosages and binary phenotypes in Phase 1, and linear regression for continuous phenotypes in phase 2. For phase 1 phenotypes, the association tests were adjusted for sex, batch, and the first four principal components, with age also included as a covariation in the AD-PGC cohort. For phase 2 (UKBB) data, age, sex, batch, and assessment centre were included as covariates. Summary statistics for 13,367,301 autosomal SNPs from Phase 3 of the analyses described in reference [1] (N samples=455,258) were made available by the Complex Trait Genetics Laboratory at VU University and VU Medical Centre, Amsterdam and were utilized in our study.

### Identification of gene expression modules

Fully processed, filtered and normalised gene expression data for 48 tissues were downloaded from the Genotype-Tissue Expression (GTEx) project portal (http://www.gtexportal.org) (version 7). Only genes with ten or more donors with expression estimates > 0.1 Reads Per Kilobase of transcript (RPKM) and an aligned read count of six or more within each tissue were considered significantly expressed. Within each tissue, the distribution of RPKMs in each sample was quantile-transformed using the average empirical distribution observed across all samples. Expression measurements for each gene in each tissue were subsequently transformed to the quantiles of the standard normal distribution.

Gene co-expression modules were individually constructed for 48 tissues (Table S1), including 13 brain tissues, using the weighted gene co-expression network analysis (WGCNA) package in R [7]. An unsigned pairwise correlation matrix – using Pearson’s product moment correlation coefficient – was calculated. An appropriate “soft-thresholding” value, which emphasises strong gene-gene correlations at the expense of weak correlations, was selected for each tissue by plotting the strength of correlation against a series (range 2 to 20) of soft threshold powers. The correlation matrix was subsequently transformed into an adjacency matrix, where nodes correspond to genes and edges to the connection strength between genes. Each adjacency matrix was normalised using a topological overlap function. Hierarchical clustering was performed using average linkage, with one minus the topological overlap matrix as the distance measure. The hierarchical cluster tree was cut into gene modules using the dynamic tree cut algorithm [11], with a minimum module size of 50 genes. We amalgamated modules if the correlation between their eigengenes – defined as the first principal component of their genes’ expression values – was greater or equal to 0.8.

### Gene-set analysis of gene co-expression modules

We performed gene-based analysis using MAGMA v1.06 [12] to (i) identify risk genes associated with AD; and (ii) test the enrichment of AD risk genes in gene co-expression modules using gene-set analysis. To identify risk genes, MAGMA assigns SNPs to their nearest gene using a pre-defined genomic window (35 kb upstream or 10 kb downstream of a gene body) and computes a gene-based statistic based on the sum of the assigned SNP – log(10) *P* values while accounting for the correlation (i.e. linkage disequilibrium) between nearby SNPs. To identify tissue-specific modules enriched with AD risk genes, we performed gene-set analysis in MAGMA. The competitive analysis tests whether the genes in a gene-set (i.e. gene co-expression module) are more enriched with Alzheimer’s disease risk genes than other genes while accounting for gene size and gene density. An adaptive permutation procedure (N=10,000 permutations) was used to obtain P values corrected for multiple testing (FDR<0.05). The 1000 Genomes European reference panel (Phase 3) was used to account for Linkage Disequilibrium (LD) between SNPs. For each tissue, a quantile-quantile plot of observed versus expected *P* values was generated to assess inflation in the test statistic.

### Biological characterisation of AD-associated gene expression modules

Gene expression modules enriched with Alzheimer’s disease GWAS association signals were assessed for biological pathways and processes using g:Profiler (https://biit.cs.ut.ee/gprofiler/) [13]. Ensembl gene identifiers within enriched gene modules were used as input; we tested for the over-representation of module genes in Gene Ontology (GO) biological processes. The g:Profiler algorithm uses a Fisher’s one-tailed test for gene pathway enrichment; the smaller the *P* value, the lower the probability a gene belongs to both a co-expression module and a biological term or pathway purely by chance. Multiple testing correction was done using g:SCS; this approach accounts for the correlated structure of GO terms, and corresponds to an experiment-wide threshold of α=0.05.

### Preservation of gene co-expression networks across tissues

To examine the tissue-specificity of biological pathways, we assessed the preservation (i.e. replication) of network modules across 48 GTEx tissues using the “modulePreservation” R function implemented in WGCNA [14]. Briefly, the module preservation approach takes as input “reference” and “test” network modules and calculates statistics for three preservation classes: i) density-based statistics, which assess the similarity of gene-gene connectivity patterns between a reference network module and a test network module; ii) separability-based statistics, which examine whether test network modules remain distinct in reference network modules; and iii) connectivity-based statistics, which are based on the similarity of connectivity patterns between genes in the reference and test networks. For simplicity, we report two density and connectivity composite statistics: “Zsummary” and “medianRank”. A Zsummary value greater than 10 suggests there is strong evidence a module is preserved between the reference and test network modules, while a value between 2 and 10 indicates weak to moderate preservation and a value less than 2 indicates no preservation. The median rank statistic ranks the observed preservation statistics; modules with lower median rank tend to exhibit stronger preservation than modules with higher median rank.

### Computational drug repurposing

Our computational drug re-purposing analysis tests the predicted effect of a drug compound on dysregulated gene expression modules underlying AD. We used S-PrediXcan to estimate the magnitude and direction of gene expression changes associated with AD. This approach integrates eQTL information with GWAS summary statistics to estimate the effect of genetic variation underlying a disease or trait on gene expression. We used eQTL information (expression weights) from 48 tissues generated by the GTEx project (v7) [15], and LD information from the 1000 Genomes Project Phase 3 [16]. These data were processed with beta values and standard errors from the GWAS of Alzheimer’s disease [1] to estimate the expression-GWAS association statistic. For each GTEx tissue, we extracted the S-PrediXcan Z-scores for genes within modules enriched with AD association signals and created two lists containing genes with either up-regulated or down-regulated expression. The gene lists were used as the basis of drug repurposing using drug gene-signatures from the Connectivity Map (CMAP) [9]. For each gene list, and for each unique compound in CMAP, we calculated a “connectivity score” based on a modified Kolmogorov-Smirnov score, which summarises the transcriptional relationship to the AD module genes. The connectivity score is a standardised metric ranging from -100 to 100, where a highly negative score indicates predicted expression effect from S-PrediXcan and the drug-gene signatures are opposing (i.e. genes that are up-regulated in disease cases are down-regulated by the compound, and vice versa). We selected compounds with connectivity scores of -90 and lower (‘strong scores’), which were subsequently mapped to mechanism of action (MOA) categories to identify chemogenomic trends. To assess the disease specificity of the CMAP enrichments, we performed a gene-based analysis of the gene co-expression networks using GWAS summary statistics for schizophrenia, a brain-related neuropsychiatric disorder with an immune component. The gene-based results were used as input to CMAP, and the results were compared with AD.

To test the significance of the AD perturbational enrichments (i.e. ensuring that significant results are not due to random chance), we grouped the observed co-expression values for pairs of genes from a single tissue (amygdala) into 100 bins based on co-expression values. We randomly sampled genes across bins, selecting the same number of gene co-expression values from each bin as the observed data. This stratified method of sampling was done to ensure the observed and permuted data were matched on connectivity. The permuted co-expression modules were uploaded to CMAP using the clue API client, and connectivity scores for each compound were extracted from the output files. We calculated empirical *P* values for observed compounds with connectivity scores smaller than -90 by counting the number of times the same compound from the permuted data had a connectivity score smaller than -90. To assess the disease specificity of the drug-gene enrichments, we ran our methodological framework using GWAS summary statistics for schizophrenia [17] and compared the overlap of significant (connectivity score ≤-90) drug-gene candidates with Alzheimer’s disease using the hypergeometric overlap test.

## Results

### Alzheimer’s disease risk genes are enriched in gene co-expression modules associated with the immune system

We used MAGMA to collapse GWAS association signals to individual genes and identified 74 genes significantly associated with AD after multiple testing correction (*P*<2.78×10^−6^) (Table S2). We tested for the enrichment of risk genes in gene co-expression modules built from 48 GTEx tissues [8]. A single gene module in 36 tissues, including 13 brain tissues, was enriched with AD risk genes (“AD modules”) (Table 1). No enrichment of gene-based association signals was observed for modules identified in whole blood, despite the larger sample size compared to many other of the tested tissues. There was moderate overlap between genes within AD modules, with 59 percent of genes specific to a single module, and only one gene (*PIK3R5*) being common to all AD modules (Figure S1). Despite this moderate gene overlap between modules, gene pathway analyses found the enrichment of immune system pathways (e.g. “immune system process” in brain amygdala, *P*=2.48 × 10^−73^, and “immune response” in brain substantia nigra; *P*=2.48 × 10^−88^) within all AD modules, including all 13 brain tissues and 23 peripheral tissues (Table 1).

**Table 1:** Gene-set enrichment analysis and biological pathway analysis of Alzheimer’s disease modules across 36 tissues in GTEx.

| Tissue | Gene set enrichment analysis |  |  |  |  | Biological pathway analysis |  |  |
| --- | --- | --- | --- | --- | --- | --- | --- | --- |
|  | N genes | Beta | SE | P | P adjust | Term | Term name | P adjust |
| Brain Amygdala | 630 | 0.157 | 0.0351 | 3.80E-06 | 0.00E+00 | GO:0002376 | immune system process | 2.48E-73 |
| Brain Anterior cingulate cortex BA24 | 353 | 0.164 | 0.047 | 2.34E-04 | 4.50E-03 | GO:0006955 | immune response | 4.54E-78 |
| Brain Caudate basal ganglia | 393 | 0.143 | 0.0442 | 6.28E-04 | 1.01E-02 | GO:0002376 | immune system process | 7.84E-73 |
| Brain Cerebellar Hemisphere | 197 | 0.248 | 0.0635 | 4.78E-05 | 7.00E-04 | GO:0006955 | immune response | 2.08E-50 |
| Brain Cerebellum | 103 | 0.425 | 0.0868 | 4.78E-07 | 0.00E+00 | GO:0002376 | immune system process | 1.60E-32 |
| Brain Cortex | 233 | 0.226 | 0.0567 | 3.32E-05 | 3.00E-04 | GO:0006955 | immune response | 1.64E-67 |
| Brain Frontal Cortex BA9 | 420 | 0.13 | 0.0423 | 1.02E-03 | 1.90E-02 | GO:0006955 | immune response | 1.14E-76 |
| Brain Hippocampus | 485 | 0.212 | 0.0395 | 3.98E-08 | 0.00E+00 | GO:0002376 | immune system process | 1.12E-95 |
| Brain Hypothalamus | 768 | 0.191 | 0.0315 | 7.53E-10 | 0.00E+00 | GO:0006955 | immune response | 1.83E-96 |
| Brain Nucleus accumbens basal ganglia | 463 | 0.171 | 0.0411 | 1.59E-05 | 4.00E-04 | GO:0006955 | immune response | 9.69E-81 |
| Brain Putamen basal ganglia | 352 | 0.175 | 0.0469 | 9.61E-05 | 1.40E-03 | GO:0006955 | immune response | 5.95E-72 |
| Brain Spinal cord cervical c-1 | 1205 | 0.0915 | 0.0248 | 1.10E-04 | 1.70E-03 | GO:0002376 | immune system process | 5.86E-77 |
| Brain Substantia nigra | 888 | 0.133 | 0.0291 | 2.44E-06 | 0.00E+00 | GO:0002376 | immune system process | 3.14E-88 |
| Adipose Subcutaneous | 638 | 0.104 | 0.0345 | 1.33E-03 | 3.50E-02 | GO:0006955 | immune response | 1.60E-76 |
| Adrenal Gland | 293 | 0.136 | 0.0502 | 3.35E-03 | 4.17E-02 | GO:0006955 | immune response | 2.34E-70 |
| Artery Aorta | 1102 | 0.0847 | 0.0266 | 7.42E-04 | 1.16E-02 | GO:0006955 | immune response | 8.41E-76 |
| Artery Coronary | 1344 | 0.0794 | 0.0243 | 5.46E-04 | 7.30E-03 | GO:0006955 | immune response | 4.75E-151 |
| Artery Tibial | 472 | 0.181 | 0.0408 | 4.95E-06 | 0.00E+00 | GO:0006955 | immune response | 4.41E-124 |
| Colon Transverse | 445 | 0.12 | 0.0412 | 1.85E-03 | 1.19E-02 | GO:0002376 | immune system process | 4.38E-91 |
| Esophagus Mucosa | 803 | 0.132 | 0.0321 | 1.88E-05 | 2.00E-04 | GO:0006955 | immune response | 1.21E-143 |
| Esophagus Muscularis | 495 | 0.122 | 0.0394 | 9.62E-04 | 1.72E-02 | GO:0006955 | immune response | 9.18E-83 |
| Heart Atrial Appendage | 371 | 0.172 | 0.0448 | 5.87E-05 | 1.60E-03 | GO:0002376 | immune system process | 4.86E-95 |
| Heart Left Ventricle | 180 | 0.275 | 0.0654 | 1.29E-05 | 2.00E-04 | GO:0006955 | immune response | 2.50E-69 |
| Liver | 611 | 0.0956 | 0.0357 | 3.72E-03 | 4.86E-02 | GO:0002376 | immune system process | 1.21E-31 |
| Lung | 146 | 0.271 | 0.0707 | 6.28E-05 | 1.20E-03 | GO:0042119 | neutrophil activation | 2.49E-36 |
| Minor Salivary Gland | 510 | 0.163 | 0.0383 | 1.04E-05 | 2.00E-04 | GO:0006955 | immune response | 1.20E-76 |
| Muscle Skeletal | 233 | 0.211 | 0.0592 | 1.81E-04 | 2.70E-03 | GO:0006955 | immune response | 6.63E-68 |
| Nerve Tibial | 283 | 0.163 | 0.0516 | 8.01E-04 | 2.49E-02 | GO:0006955 | immune response | 6.78E-66 |
| Pancreas | 852 | 0.151 | 0.0303 | 3.11E-07 | 0.00E+00 | GO:0006955 | immune response | 7.78E-113 |
| Skin Not Sun Exposed Suprapubic | 178 | 0.213 | 0.0681 | 8.84E-04 | 1.21E-02 | GO:0002376 | immune system process | 1.46E-59 |
| Small Intestine Terminal Ileum | 2327 | 0.0506 | 0.0181 | 2.65E-03 | 2.72E-02 | GO:0002376 | immune system process | 6.74E-39 |
| Spleen | 963 | 0.0878 | 0.0279 | 8.18E-04 | 2.44E-02 | GO:0042119 | neutrophil activation | 3.42E-45 |
| Stomach | 72 | 0.339 | 0.104 | 5.59E-04 | 5.80E-03 | GO:0006955 | immune response | 1.87E-17 |
| Thyroid | 979 | 0.0879 | 0.0292 | 1.28E-03 | 3.49E-02 | GO:0002376 | immune system process | 4.10E-114 |
| Uterus | 355 | 0.198 | 0.0472 | 1.33E-05 | 4.00E-04 | GO:0006955 | immune response | 4.67E-85 |
| Vagina | 149 | 0.269 | 0.0728 | 1.12E-04 | 5.00E-04 | GO:0006955 | immune response | 1.01E-47 |

### Gene co-expression modules enriched with Alzheimer’s disease risk genes are preserved across brain tissues

We assessed the preservation (i.e. reproducibility) of AD modules across 36 enriched tissues using the WGCNA *modulePreservation* algorithm. Strong modular preservation (z-score ≥ 10) was observed across most tissues; preservation tended to be higher between vascular-related tissues (e.g. coronary artery, aorta, and tibial artery) and all other tissues, and among brain-related tissues. The weakest preservation (z-score < 10) of immune modules was observed in stomach, vagina, spleen, and lung tissues (Figure 1). These data suggest immune changes in AD are systematic (that is, may be detected in brain and peripheral tissues), but are likely greater (or manifest earlier) in brain tissues. Therefore, the study of gene co-expression—that is, the connectivity between genes—across multiple brain and peripheral tissues, such as skin, may provide a useful surrogate for genetic and molecular studies of brain-related processes in AD.

### A computational drug re-purposing analysis identifies drug compounds for further analysis

Our gene co-expression analytical approach is built on the premise that genes do not act alone, but rather form complex networks to influence the manifestation of a disease or trait [18]. A similar premise can be applied to therapeutics, where drugs are likely to not only modulate the activity of a single gene but the activity of multiple, highly connected genes in a pathogenic tissue. Our multi-tissue gene co-expression modules provide a useful substrate for the identification and prioritisation of drugs that may “normalise” altered gene co-expression in AD. We used S-PrediXcan to identify genes whose expression is associated with genetic variation underlying AD (Table S3). We assigned the S-PrediXcan Z-score for the direction and magnitude of effect to all genes within AD modules, and generated lists of up-regulated and down-regulated genes. The gene lists were used as input to the connectivity map (CMAP), which computes a “connectivity score” based on the transcriptional relationship between the gene lists and observed drug-gene signatures across multiple cell-types. We selected compounds with connectivity scores less than or equal to -90 (indicating a compound is predicted to normalise predicted altered [up- or down-regulated] expression patterns in AD) (Table S4). We collapsed significant compounds based on their mechanism of action across cell types and brain tissues (Table S5). Top-ranked mechanisms of action included acetylcholine receptor antagonists, which include a number of currently-used drugs for AD (e.g. memantine). Other top-ranked mechanisms included GABA receptor modulators, which are reported to have a key role in autoimmune inflammation in the brain [19]. Other mechanisms of action included cyclooxygenase inhibitors, which include drug compounds that may have a protective role in Alzheimer’s disease [20]. In order to assess the significance of drug-gene level results, we applied a permutation procedure (methods) in Amygdala. The results show top-ranked compounds are unlikely to be due to correlated expression (Table S6). We also ran our network-based pipeline with GWAS summary statistics for schizophrenia, a brain-related trait with an immune component, and found no significant overlap (hypergeometric test) with our observed results for Alzheimer’s disease across cell types (Table S7). These observations strengthen the candidacy of potential AD therapeutics and illustrate the potential of CMAP within a gene co-expression network framework to generate novel, unbiased hypotheses on the pharmacologic modulation of disease states.

## Discussion

We applied a tissue-specific network-based gene co-expression method to identify groups of highly correlated (and functionally related) genes associated with AD. Gene-based analyses of GWAS summary statistics identified 74 genes significantly associated with AD after multiple testing correction. Gene-based associations were enriched in a single gene module in 36 GTEx tissues, including 13 brain tissues. All tissue-specific AD modules were associated with the immune system and immune response, despite only moderate overlap in gene membership across modules. A computational drug repositioning analysis of genes within AD modules identified the mechanism of action categories acetylcholine receptor antagonists and glutamate receptor function, both of which include drug candidates, in addition to cyclooxygenase inhibitors and other immune function-related drug classes. Our results demonstrate a tissue-specific approach to gene discovery in AD may not only identify candidate causal genes, tissues, and biological pathways, but also targets for therapeutic intervention.

Many studies suggest an important role of neuroinflammation in the onset and progression of the neuropathological changes that are observed in AD. Independent studies had identified immune-related proteins and cells in the proximity of β-Amyloid plaques [21], for example, and epidemiological reports suggested anti-inflammatory agents used to treat immune disorders, such as rheumatoid arthritis, decrease the risk of AD [22]. It was not until the publication of a large-scale GWAS on AD that the first robust evidence for a causal association between neuroinflammation and disease onset could be established [1]. However, the GWAS only found enrichment of immune-related tissues and cell types, rather than a biological mechanism. The study of tissue-specific gene co-expression patterns allowed us to investigate a larger set of genes that might be implicated in disease based on network connectivity. Using this approach, we identified immune system-related tissue-specific modules (groups) of co-expressed genes that are both enriched with AD association signals and strongly preserved across tissues.

We further demonstrate the versatility of co-expression network-based methods with the application of a computational drug repositioning analysis. We used all genes within disease implicated co-expression modules as input to CMap, under the biologically valid assumption that a drug compound not only alters the activity of a single target gene, but influences the activity of multiple related genes through co-regulation [23]. Furthermore, by using S-PrediXcan to impute expression effects from GWAS summary statistics, we focus only genetically regulated gene expression effects, thereby removing unwanted variation on gene expression from environmental effects. This approach identified disease-associated gene modules whose expression signature was predicted to be normalised by major Alzheimers’s disease drug classes, including acetylcholine receptor antagonists and glutamate receptor antagonists. We also found several drug classes that target immune-related processes,, consistent with current knowledge on disease pathways underlying AD. For example, proliferator-activated receptor γ (PPAR) agonists were predicted to normalise gene expression signatures in AD. PPARs play an important role in microglial-driven inflammatory response, and their activation is known to increase microglial amyloid β clearance and improve spatial memory performance in a mouse model [24]. Other mechanism of action classes, such a cyclooxygenase inhibitors, contained compounds that have undergone clinical trials for AD (e.g. naproxen) [25]. Collectively, these results show our approach can uncover existing and potentially novel drugs for further functional studies in appropriate cell types and model organisms.

Genetic associations for AD are enriched in genomic regions that encode “druggable” gene targets [26] and therefore has translational potential using computational repositioning methods. So *et al.* [27] used GWAS-imputed transcriptome profiles and the CMAP algorithm to identify candidates for drug repositioning in neuropsychiatric disorders and identified several non-steroidal anti-inflammatory drugs (NSAIDs) with possible benefits in AD, in line with our results. We extend this analysis with the use of a larger, more highly powered GWAS and network-based methods to implicate additional genes with potential relevance in AD pathology. A recently published study used genetic information and network-based methods to develop a priority index for drug target validation in immune-mediated traits [28]. The priority index incorporated functional genomic information with protein-protein network connectivity information, and was shown to successfully identify current therapeutics and prioritise alternative compounds for early-stage testing. Their network annotations, however, do not directly integrate genetic co-expression, but instead rely on disparate sources of protein interaction data to characterise gene connectivity. Our gene co-expression-based approach, on the other hand, directly anchors changes in genetically regulated gene expression to observed levels of co-expression between genes, and therefore more closely represents underlying biological relationships.

We used gene expression data from bulk human brain tissue as single cell expression data is not available in GTEx. Bulk brain tissue is not homogeneous with respect to individual cell types (e.g. microglial cells versus neurons). As a result, true Alzheimer’s disease association signals may be diluted by non-specific expression, or expression differences may simply reflect mosaic effects of different cell types. This is especially problematic for AD, where many of the risk genes are not highly expressed in bulk brain tissues [29,30] and have been linked to immune function [31]. Genetic signals for AD are enriched in microglial cells [32], which themselves only account for around 3% of the total brain cell population. Therefore, RNA sequencing of individualised cells (known as single cell RNA sequencing) may partition genetic signals to causal cell types and improve power to identify functional genes and mechanisms underlying AD and, in turn, improve the accuracy of drug positioning [6]. An alternative approach is to use human monocyte-induced microglia as a proxy to examine the genetic effects of AD in microglia. This approach would remove the need for post-mortem brain samples, which can be difficult to obtain, and thereby enable the rapid functional validation of molecular process and drug responses in AD.

In summary, we integrated GWAS summary statistics for AD with gene co-expression networks from 48 tissues in GTEx. We identified gene modules within each of the tissue-specific co-expression networks enriched with AD GWAS association signals. Biological pathway analysis of the implicated gene modules found enrichment of immune-response pathways, consistent with recent functional and transcriptomic analyses [5]. We estimated the effect direction of gene expression underlying AD using S-PrediXcan and integrated these data with drug-gene signatures from the connectivity map. This procedure identified drug compounds whose effect on gene expression was estimated to “normalise” dysregulated expression patterns underlying AD. We collapsed drugs to common mechanisms of action, and found major drug classes in the treatment (glutamate receptor antagonists and acetylcholinesterase inhibitors) were captured by our method, in addition to other mechanisms of action of biological interest (e.g. cyclooxygenase). We identified a list of drug compounds with genetic support that may be repositioned for the treatment of AD after follow-up functional studies. Our approach will help researchers to translate genetic findings for follow-up functional studies in the early-state drug development process.

---

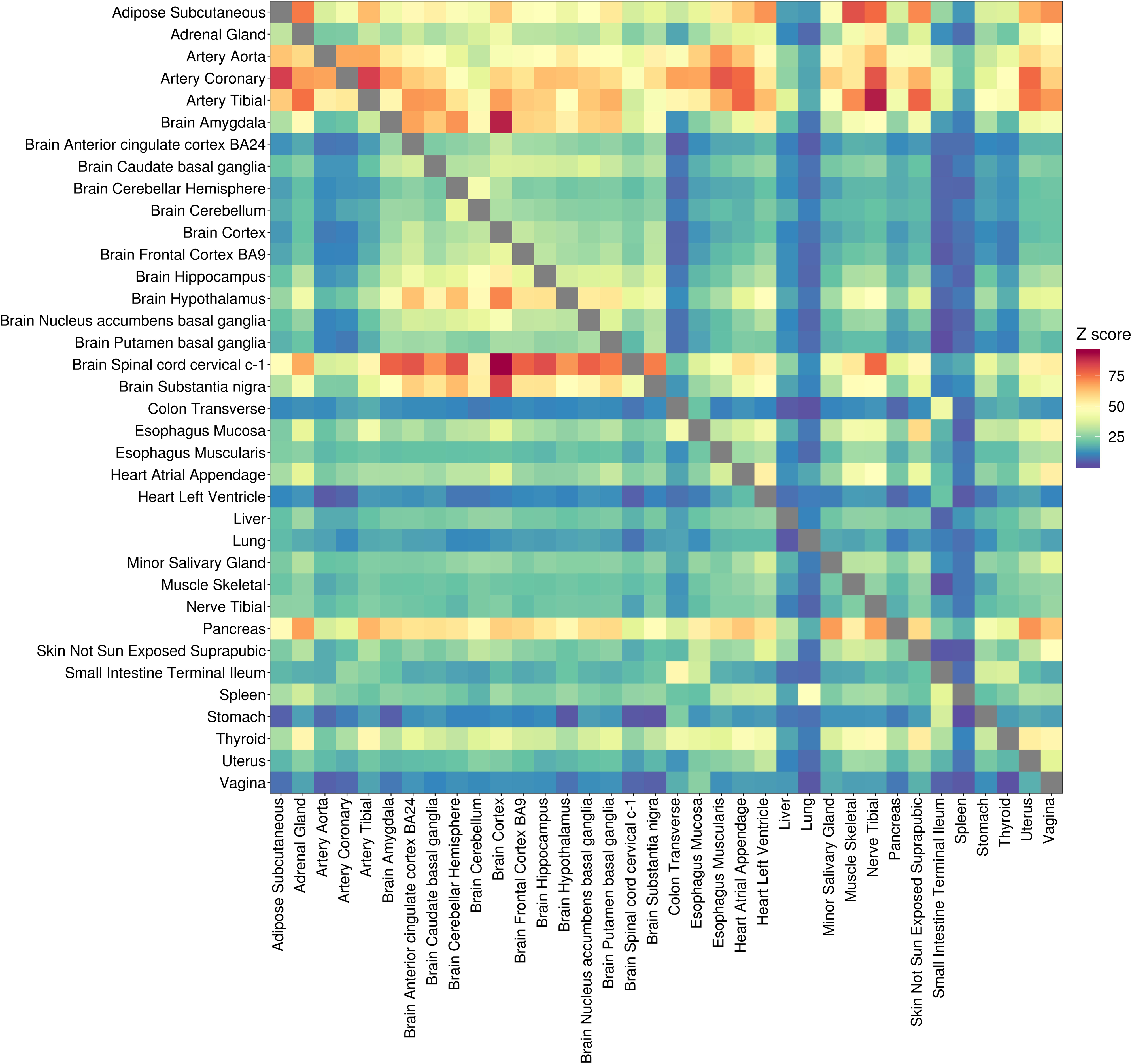

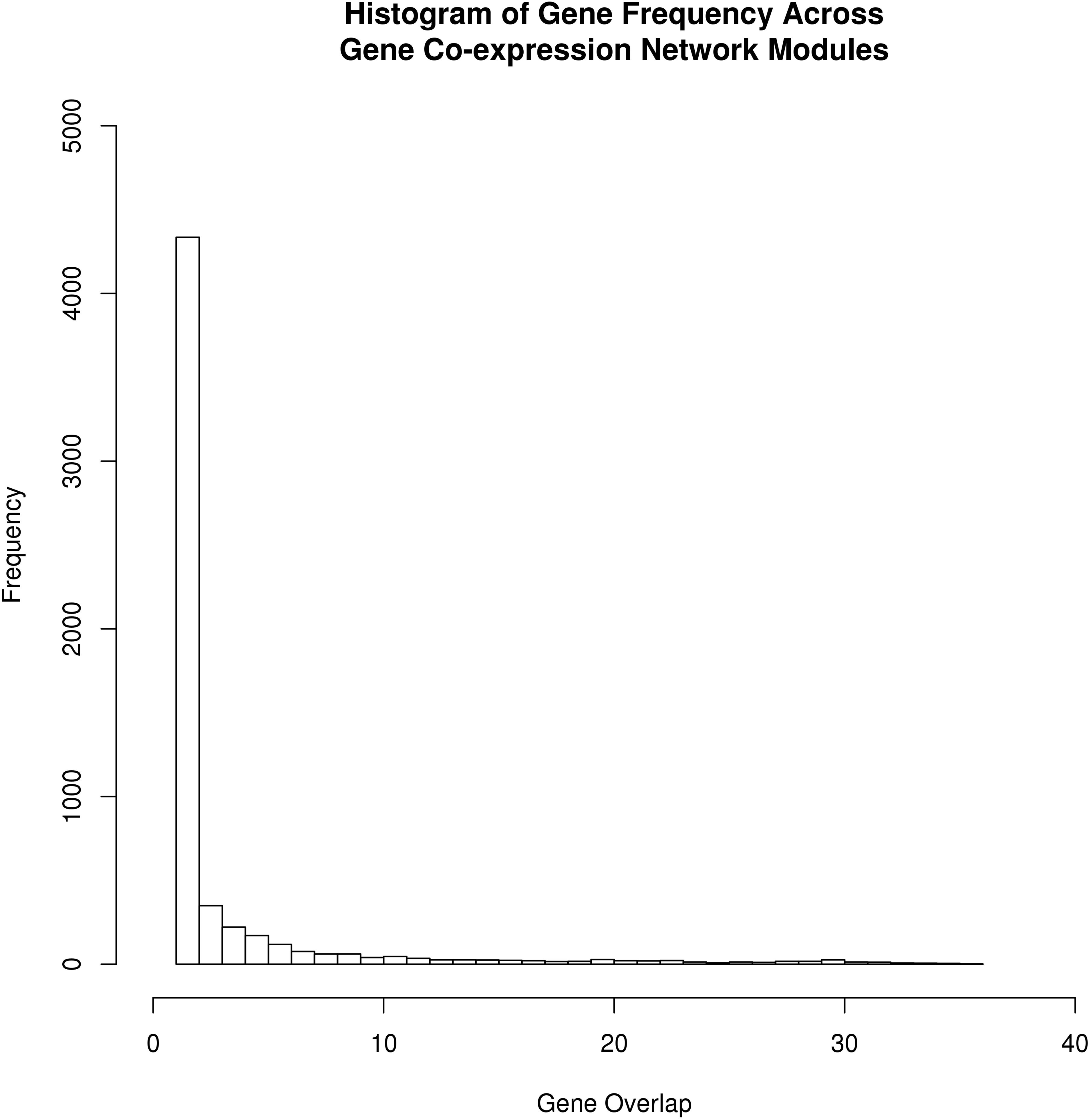

